# Efficient flavinylation of glycosomal fumarate reductase by its own ApbE domain in *Trypanosoma brucei*

**DOI:** 10.1101/2020.12.30.424822

**Authors:** Robin Schenk, Sabine Bachmaier, Frédéric Bringaud, Michael Boshart

**Author notes:** **Correspondence**: Prof. Michael Boshart, Biozentrum, Fakultät für Biologie, Genetik, Ludwig-Maximilians-Universität München (LMU), 82152, Martinsried, Germany. Biozentrum, Fakultät für Biologie, Genetik, Ludwig-Maximilians-Universität München (LMU), 82152, Martinsried, Germany. CNRS, Microbiologie Fondamentale et Pathogénicité (MFP), UMR 5234, Université de Bordeaux, F-33000, Bordeaux, France.

## Abstract

A subset of flavoproteins has a covalently attached flavin prosthetic group enzymatically attached via phosphoester bonding. In prokaryotes, this is catalysed by ApbE flavin transferases. ApbE-like domains are present in few eukaryotic taxa, e.g. the N-terminal domain of fumarate reductase (FRD) of *Trypanosoma*, a parasitic protist known as a tropical pathogen causing African sleeping sickness. We use the versatile reverse genetic tools available for *Trypanosoma* to investigate the flavinylation of glycosomal FRD (FRDg) *in vivo* in the physiological and organellar context. Using direct in-gel fluorescence detection of covalently attached flavin as proxy for activity, we show that the ApbE-like domain of FRDg has flavin transferase activity *in vivo*. The ApbE domain is preceded by a consensus flavinylation target motif at the extreme N-terminus of FRDg, and serine 9 in this motif is essential as flavin acceptor. The preferred mode of flavinylation in the glycosome was addressed by stoichiometric expression and comparison of native and catalytically dead ApbE domains. In addition to the *trans*-flavinylation activity, the ApbE domain catalyses the intramolecular *cis*-flavinylation with at least 5-fold higher efficiency. We discuss how the higher efficiency due to unusual fusion of the ApbE domain to its substrate protein FRD may provide a selective advantage by faster FRD biogenesis during rapid metabolic adaptation of trypanosomes. The first 37 amino acids of FRDg, including the consensus motif, are sufficient as flavinylation target upon fusion to other proteins. We propose FRDg(1-37) as 4 kDa heat-stable, detergent-resistant fluorescent protein tag and suggest its use as a new tool to study glycosomal protein import.

## 4 Introduction

Flavoproteins are ubiquitous redox-proteins associated with an eponymous flavin molecule, most commonly, flavin adenine dinucleotide (FAD) or its precursor, flavin mononucleotide (FMN). A defining feature shared by those cofactors is the isoalloxazine ring (Figure S1), a tricyclic structure, which introduces a wide variety of chemical and physical properties, in particular when incorporated into the active centre of enzymes. Versatility of catalytic reactions is exemplified by numerous oxidases, reductases, dehydrogenases, electron transferases, mediating both one- and two-electron transfer. Flavoproteins also include DNA damage repair proteins and photochemical signalling components [1–4]. Blue-light photoreceptors and bacterial luciferases [5, 6] utilise the distinct fluorescent properties of flavins for their function [7]. They typically feature excitation/emission maxima of the oxidised form at around 450/520 nm, depending on environmental conditions and binding partners [8, 9]. The optical properties do not only allow characterisation of specific flavoproteins and their state transitions [10, 11], but also visualisation of flavin-rich cell populations or subcellular structures such as mitochondria [12, 13].

Flavin cofactors are mostly tight binding and structurally shielded, presumably to prevent autoxidation [2]. Yet, non-covalent binding strongly prevails, with only ~10 % of flavoproteins being covalently flavinylated. This minority of enzymes has an increased redox potential allowing for thermodynamically demanding reactions and more efficient substrate turnover, as well as higher holoenzyme stability due to fixation of the cofactor [14, 15]. Mechanistically, linkage of cysteine, tyrosine or histidine to the 8*α*-methyl group or the C6 atom of the isoalloxazine ring (Figure S1) has been reported and suggested to occur auto-catalytically [16]. Alternatively, in prokaryotes flavin transferases of the ApbE (alternative pyrimidine biosynthesis; also Ftp: flavin-trafficking protein) family (PFAM accession: PF02424) attach FMN covalently to threonine residues by phosphoester bond formation [17, 18]. ApbE recognises the degenerated consensus sequence Dxx(s/t)gA<u>T</u> and targets threonine 7 in this motif as the FMN acceptor residue [19, 20]. A well described target of ApbE-mediated post-translational modification is Na^+^-translocating NADH:quinone oxidoreductase, a component of the respiratory chain [17, 21].

Fumarate reductases (FRDs), which catalyse the reduction of fumarate to succinate during anaerobic metabolism [22, 23], predominantly occur as multi-subunit membrane-bound complexes. In bacteria, the FRD periphery is exposed to the cytoplasm, in eukaryotes to the mitochondrial matrix, transferring electrons from quinol to the terminal electron acceptor fumarate. In some organisms, a second type of FRD is found that is soluble and monomeric, oxidises FADH_2_, FMNH_2_ or NADH in place of quinol [24] and contains a cytochrome domain instead of an iron sulphur cluster subunit [25]. Both classes of FRD further contain a prosthetic flavin group [22]. While evidence points to auto-catalytic linkage of flavin to membrane-bound FRD in *E. coli* [26, 27], in *Klebsiella pneumoniae* presence of ApbE is essential for activity of its cytoplasmic fumarate reductase [19, 28], suggesting a covalent attachment *in trans* by the ApbE protein. In this context it is particularly interesting that the soluble NADH-dependent FRD in kinetoplastids [24, 29], an evolutionary distant branch of eukaryotes, contains an ApbE-like domain at the N-terminus of the protein. *Kinetoplastida* include pathogenic parasite species that cause major tropical diseases, such as sleeping sickness, chagas disease or leishmaniasis. The ApbE domain is widespread in bacteria and archaea, yet only present in very few eukaryotes, notably the kinetoplastids. The model kinetoplastid *Trypanosoma brucei* encodes three NADH-dependent FRD isoforms. FRDg is localised in glycosomes, specialised peroxisome organelles harbouring glycolytic enzymes in kinetoplastids [30]. FRDm1 is localised in the mitochondrion, whereas for *FRDm2* no expression or enzymatic activity was detected so far [29]. FRDg and FRDm1 are essential for a metabolic process termed succinic fermentation in the glycosomes and the mitochondrion [24]. Despite their role in maintaining the redox equilibrium by oxidation of NADH to NAD^+^, none of the isoforms alone is essential for growth of the parasite in culture conditions [24, 29, 31].

The predicted ApbE domain of kinetoplastid FRDs is preceded by a putative flavinylation target motif (FTM; Figure 1A, B) with a Dxx(s/t)(s/g)AS consensus sequence that slightly diverges from the prokaryotic consensus [32]. For the trypanosomatid *Leptomonas pyrrhocoris*, covalent flavin attachment at serine 9 of FRDg within the N-terminal FTM was verified by mass spectrometry [32]. Replacement of S9 by an asparagine residue abolished both flavinylation and NADH-fumarate reductase activity of FRDg upon expression in yeast. Therefore, it was suggested that the ApbE-like domain may catalyse the transfer of FMN from FAD to serine 9 of *L. pyrrhocoris* FRDg [32].

**Figure 1:**
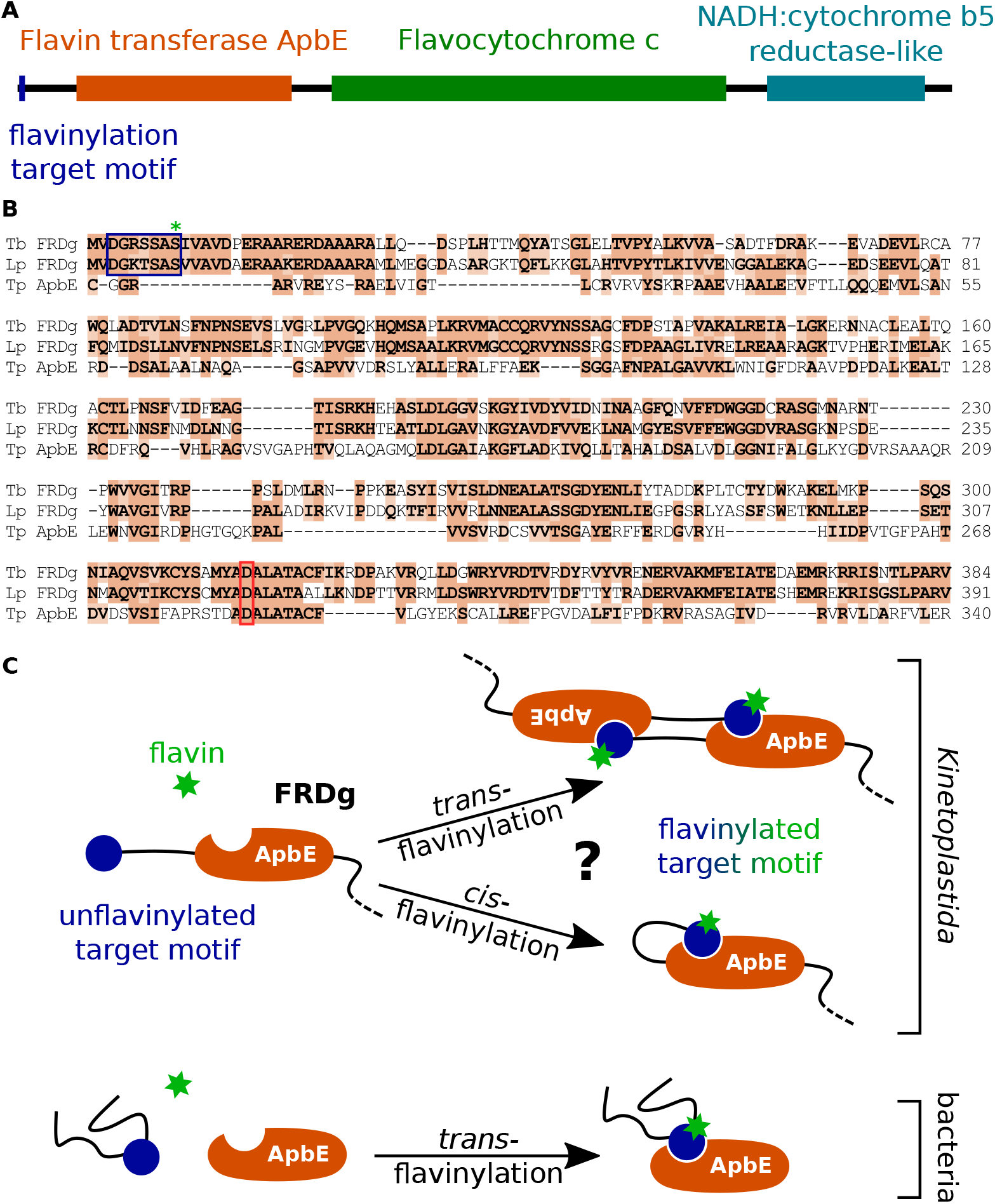
Domain overview of kinetoplastid FRDg and sequence comparison with bacterial ApbE. **(A)** A true-to-scale representation of the *T. brucei* FRDg (TriTrypDB accession: Tb927.5.930) domain architecture as predicted by InterPro [59]. Orange box: flavin transferase ApbE domain (InterPro accession: IPR024932); green box: the flavocytochrome c domain (InterPro accession: IPR010960) that contributes the NADH:fumarate reductase catalytic centre [87]; blue box: the NADH:cytochrome b5 reductase-like domain (InterPro accession: IPR001834). **(B)** Protein sequence alignment of the *T. brucei* FRDg N-terminus, the *L. pyrrhocoris* FRDg N-terminus (Lp; TriTrypDB accession: LpyrH10 01 2930) and *Treponema pallidum* ApbE (Tp; GenBank accession: AAC65759). Similar amino acids are highlighted in orange; The flavinylation target motif (FTM) is indicated by a blue frame, a green asterisk marking the flavin accepting Ser9 in LpFRDg. The residues aligned to Mg^2+^ binding Asp284 of TpApbE are marked by a red frame [62]. **(C)** Possible modes of flavinylation in different species. A model for intermolecular *trans*- and intramolecular *cis*-flavinylation by the ApbE domain of kinetoplastid FRD, compared to *trans*-flavinylation by ApbE in bacteria [17].

Here we use the reverse genetic tools available for the model kinetoplastid *T. brucei* to show that the N-terminal ApbE-like domain of FRDg functions as flavin transferase *in vivo*. A minimal target sequence sufficient for flavinylation by the ApbE domain of FRD was defined that coincides with the consensus sequence. Alternative modes of ApbE-mediated flavinylation (Figure 1C) are addressed by domain swap experiments and provide evidence for significantly higher efficiency of intramolecular *cis*-flavinylation compared to *trans*-flavinylation activity. We suggest that fusion of an ApbE domain to FRD may provide an evolutionary advantage.

## 5 Materials and Methods

### 5.1 *T. brucei* cultivation

The bloodstream form (BSF) of AnTat1.1 90-13, a pleomorphic *T. brucei brucei* cell line constitutively expressing T7 RNA polymerase and tetracycline repressor [33, 34], was cultured at 37°C and 5 % CO_2_-atmosphere in modified HMI-9 medium [35] containing 10 % (V/V) heat inactivated fetal calf serum (FCS). The procyclic form (PCF) was cultivated at 27°C in modified SDM-79 medium [36] supplemented with 10 % (V/V) heat inactivated FCS, 7.5 *mg l*^−1^ hemin, 10 mM glucose and 10 mM glycerol. Continuous selection was performed with hygromycin B (BSF: 2.5 *μg ml*^−1^; PCF: 25 *μg ml*^−1^), G418 (BSF: 2.0 *μg ml*^−1^; PCF: 15 *μg ml*^−1^). To differentiate BSF to PCF, cultures were started at a density of 5.0 * 10^5^ *ml*^−1^. Growth was continued for 36 h without dilution. Usually, 10 ml of culture were harvested, resuspended in 5 ml modified DTM medium [35] supplemented with 6 mM cis-aconitate and continued as procyclic cultures by dilution with SDM-79 upon resuming proliferation.

### 5.2 Transgenic cell lines

To express *FRDg(1-350)*, a C-terminal truncation of FRDg (TriTrypDB accession: Tb927.5.930) with a length of 350 amino acids, followed by two Ty1 epitopes [37], linked and interspaced with (PS)_x1_ linkers (DNA: CCAAGT), was inserted between the HindIII and BamHI restriction sites of the tetracycline-inducible pLew82 [38] expression vector. The respective fragment was amplified from genomic DNA of strain AnTat1.1 Munich [34]. NotI was used for plasmid linearisation before transfection of the AnTat1.1 90-13 cell line with the *Nucleofector II* device (Amaxa Biosystems) [39]. Continuous selection was performed with phleomycin (BSF: 2.0 *μg ml*^−1^; PCF: 2.5 *μg ml*^−1^).

The same approach was used to express *FRDg(1-363)*, *FRDg(1-363)-S9N* that is identical to *FRDg(1-363)* except for a substitution of serine 9 (TCA) with asparagine (AAT) and *FRDg(1-363)-D316A* that is identical to *FRDg(1-363)* except for a substitution of aspartic acid 316 (GAC) with alanine (GCA).

FRDg(1-37) was N-terminally fused to *T. brucei* aconitase (*ACO*) (TriTrypDB accession: Tb927.10.14000) [40], replacing the mitochondrial targeting signal (FRDg(1-37)-ACO). Using XbaI, two fragments were inserted simultaneously between the HindIII and BamHI restriction sites of pLew82 with a puromycin resistance cassette; (1) a fragment containing the first 111 bp of the *FRDg* gene, followed by a flexible GSAGSAAGSGEF linker [41] (DNA: GGGTCTGCGGGGTCTGCAGCTGGGAGTGGTGAGTTT), (2) a 2664 bp fragment of *ACO* position 31-2694. AnTat1.1 90-13 and the derived FRDg(1-363) cell line were transfected with the plasmid. Continuous selection was performed with puromycin (BSF: 0.1 *μg ml*^−1^; PCF: 1.0 *μg ml*^−1^).

FRDg(1-363) was N-terminally fused to glycosomal isocitrate dehydrogenase (TriT-rypDB accession: Tb927.11.900), resulting in the fusion protein FRDg(1-363)-IDHg. A fragment containing the first 1089 bp of the *FRDg* gene, followed by a GSAGSAAGSGEF linker, was inserted between the HindIII and AgeI restriction sites of an *IDHg* open reading frame (TriTrypDB accession: Tb927.11.900) containing plasmid [42] based on the constitutive expression vector pTSARib [43]. BglII was used for plasmid linearisation before transfection of the AnTat1.1 90-13 cell line. Continuous selection was performed with puromycin (BSF: 0.1 *μg ml*^−1^; PCF: 1.0 *μg ml*^−1^).

The same approach was used to express three additional FRDg-IDHg fusion proteins: (1) *FRDg(1-9)-IDHg* (FRD(1-9) N-terminally fused to IDHg), (2) *FRDg(1-37)-IDHg* (FRD(1-37) N-terminally fused to IDHg) and (3) *FRDg(1-363)-D316A-IDHg* (FRD(1-363)-D316A N-terminally fused to IDHg).

Primer sequences are available upon request.

### 5.3 SDS-PAGE and western blot analysis

Cells were washed in PBS and resuspended to 3.0 * 10^5^ *μl*^−1^ in Laemmli sample buffer (58.33 mM Tris-HCl (pH 6.8); 200 mM DTT; 5 % (V/V) glycerol; 1.71 % (m/V) SDS; 0.002 % (m/V) bromophenol blue; solvent: H_2_O). After incubation at 95°C for 5 min the sample was sonicated 3 times for 30 sec on/off with a *Bioruptor* (*Diagenode*) on high power. 10 *μ*l of the sample was subjected to SDS-PAGE (8 % or 10 %). Western blotting was performed onto a PVDF membrane with the *Trans-Blot Turbo Transfer System* (*Bio-Rad Laboratories*). Mouse *α*-PFR-A/C (clone: L13D6; 1:2000) [44], mouse *α*-Ty1 (clone: BB2; 1:500) [37], rabbit *α*-IDHg (1:5000) [42], rabbit *α*-ACO (1:750) [45], rat *α*-ACO (1:750) [45], rabbit *α*-PPDK (1:1000) [46], mouse *α*-HSP60 (1:5000) [47, 48] and rabbit *α*-ENO (1:100,000; gift from F. Bringaud, Bordeaux, France; originating from P. Michels, Edinburgh, UK) were used as primary antibodies for immunodetection. For near-infrared fluorescence detection goat IR-BLOT 800 *α*-mouse IgG (1:5000; *Cyanagen Srl*) or goat IRDye 800CW *α*-mouse IgG (H + L) (1:5000; *LI-COR Biosience*), goat IRDye 800CW *α*-rat IgG (H + L) (1:5000; *LI-COR Biosience*) and goat IRDye 680LT *α*-rabbit IgG (H + L) (1:5000; *LI-COR Biosience*) were used as secondary antibodies. Image acquisition was performed with the *Odyssey CLx Near-Infrared Fluorescence Imaging System* (LI-COR Biosience).

### 5.4 Flavinylation analysis by in-gel fluorescence

As described before [49], SDS-PAGE gels were scanned with a *Typhoon TRIO Variable Mode Imager System* (GE Healthcare). *λ*_ex_ = 488 nm and *λ*_em_ = 526 nm were used for detection of covalently bound flavin and *λ*_ex_ = 670 nm and *λ*_em_ = 633 nm for visualisation of the *Blue Prestained Protein Standard* (New England Biolabs).

### 5.5 Subcellular digitonin fractionation

2.0 * 10^9^ procyclic *T. brucei* cells were harvested and washed twice in PBS. The cells were resuspended to 6.5 * 10^8^ *ml*^−1^ in STEN buffer (250 mM sucrose; 150 mM NaCl; 25 mM Tris-HCl (pH 7.4); 1 mM EDTA) supplemented with 3.5 % (V/V) PMSF (57 mM; solvent: isopropanol). 200 *μ*l aliquots of the cell suspension (corresponding to 660 *μ*g protein [50]) were incubated for 4 min at 25°C in 14 different digitonin concentrations. Immediately after incubation, the samples were centrifuged at 14,000 g for 2 min. The supernatant was analysed by SDS-PAGE and western blotting. Pyruvate phosphate dikinase (PPDK), heat shock protein 60 (HSP60) and enolase (ENO) were used as glycosomal, mitochondrial and cytoplasmic marker proteins, respectively. The maximum signal intensity obtained for each analysed protein was set to 1 and the data points were fitted to the logistic function *f* (*x*) = *A/*(1 + *exp*((*x* − *m*)*/s*)) by application of the Levenberg-Marquardt algorithm for non-linear least square analysis. Therefore, the Java library *least-squares-in-java* [51] was used. The initial parameter estimates were set as (*A* = 10, *m* = 250, *s* = 10). The data were further normalised to the respective inferred upper asymptote *A* for graphical representation.

### 5.6 Partial purification and protein identification

2.0 * 10^9^ procyclic *T. brucei* cells of AnTat1.1E Paris [34] were harvested and washed 2 times in PBS. The cells were resuspended in 2.4 ml hypotonic lysis buffer (5mM Tris-HCl (pH 7.4); 1.995 mM PMSF) and incubated 20 min on ice. All centrifugations were performed at 4°C. Samples were centrifuged for 15 min at 1,500 g. The supernatant was subsequently centrifuged for 15 min at 3,000 g. The pellet resulting from the last centrifugation step was washed with 500 *μl* hypotonic lysis buffer by centrifugation at 3,000g for 15 min. The supernatant was discarded. After resuspension in hypotonic lysis buffer, centrifugation at 20,000 g for 15 min was performed. The pellet was washed in 100 *μl* hypotonic lysis buffer and was resuspended in 30 *μl* Laemmli sample buffer. 15 *μl* were subjected to SDS-PAGE. The gel was stained with Colloidal Coomassie Brilliant Blue G-250 [52, 53]. The partially purified protein band was identified by in-gel fluorescence, excised (Figure S2A) and destained, followed by in-gel trypsin digestion. Isolated peptides were purified using C18 stage tips and analysed by LC-MS/MS. Proteomic analysis was performed at the Protein Analysis Unit (ZfP) of the Ludwig Maximilian University of Munich, a registered research infrastructure of the Deutsche Forschungsgemeinschaft (DFG, RI-00089). MaxQuant [54] software was used for peptide identification and quantification.

### 5.7 Bioinformatic analysis and toolchain

For statistical analysis and plotting R 3.6.2 [55] was used with the integrated development environment RStudio 1.1.463 [56]. In silico cloning and sequence alignment was performed with CLC Main Workbench 8.1 (https://www.qiagenbioinformatics.com). Sequence similarity was visualised using Sequence Manipulation Suite 2 [57]. Image post-processing was carried out with Fiji [58], quantification with Image Studio Lite (https://www.licor.com). Protein domains were annotated and predicted using InterPro [59], while general genomic information was retrieved from TriTrypDB [60] or GenBank [61].

## 6 Results

### 6.1 Flavin transferase activity of trypanosome FRDg

To trace a prominent endogenous auto-fluorescent band in *T. brucei* cell lines upon in-gel fluorescence analysis [49], we enriched the respective band by cell fractionation and identified it by mass spectrometry (Figure S2) as the flavoprotein fumarate reductase (FRD). This explained the optical properties [8, 9] and resistance of the fluorescence to denaturing conditions, as covalent flavin attachment to FRD had been reported in the closely related kinetoplastid *L. pyrrhocoris* [32]. The presence of an ApbE-like domain at the N-terminus of FRDg led to the hypothesis of covalent FMN attachment catalysed by a eukaryotic ApbE flavin transferase activity [32]. We tested this hypothesis using the technically more amenable kinetoplastid model organism *T. brucei* that shares the domain structure and a predicted flavinylation target motif Dxx(s/t)(s/g)AS with *L. pyrrhocoris*.

We first asked if the ApbE-like domain has flavin transferase activity in trypanosomes *in vivo*. The N-terminus of TbFRDg with the ApbE domain and the predicted flavinylation target motif and site-directed mutants thereof (Figure 2A) were transgenically expressed in *T. brucei* strain Antat1.1 90-13 and analysed in the procyclic developmental stage of the parasite. To exclude possible phenotypic effects due to expression of the truncated proteins, an inducible expression system based on the *E. coli tet* operon was used. Expression of the recombinant proteins was quantified by western blotting (Figure 2D) using the Ty epitope tag. Some leakiness of the inducible expression system was noticed. Covalent flavinylation was quantified by in-gel fluorescence analysis of denaturing SDS-PAGE gels. This was possible since the only prominent auto-fluorescent bands in wild type cells are the fumarate reductases FRDg and FRDm1 (Figure S3A-F). The expressed ApbE domain lacks the C-terminal glycosomal targeting signal of FRDg, resulting in cytoplasmic localisation of the truncated protein (Figure 2B). The endogenous FRDg is glycosomal and FRDm1 is mitochondrial, due to their C-terminal and N-terminal topogenic signals, respectively [29]. Therefore, we excluded significant flavinylation of the transgenically expressed truncated proteins by the endogenous FRD paralogues that both contain predicted ApbE domains.

**Figure 2:**
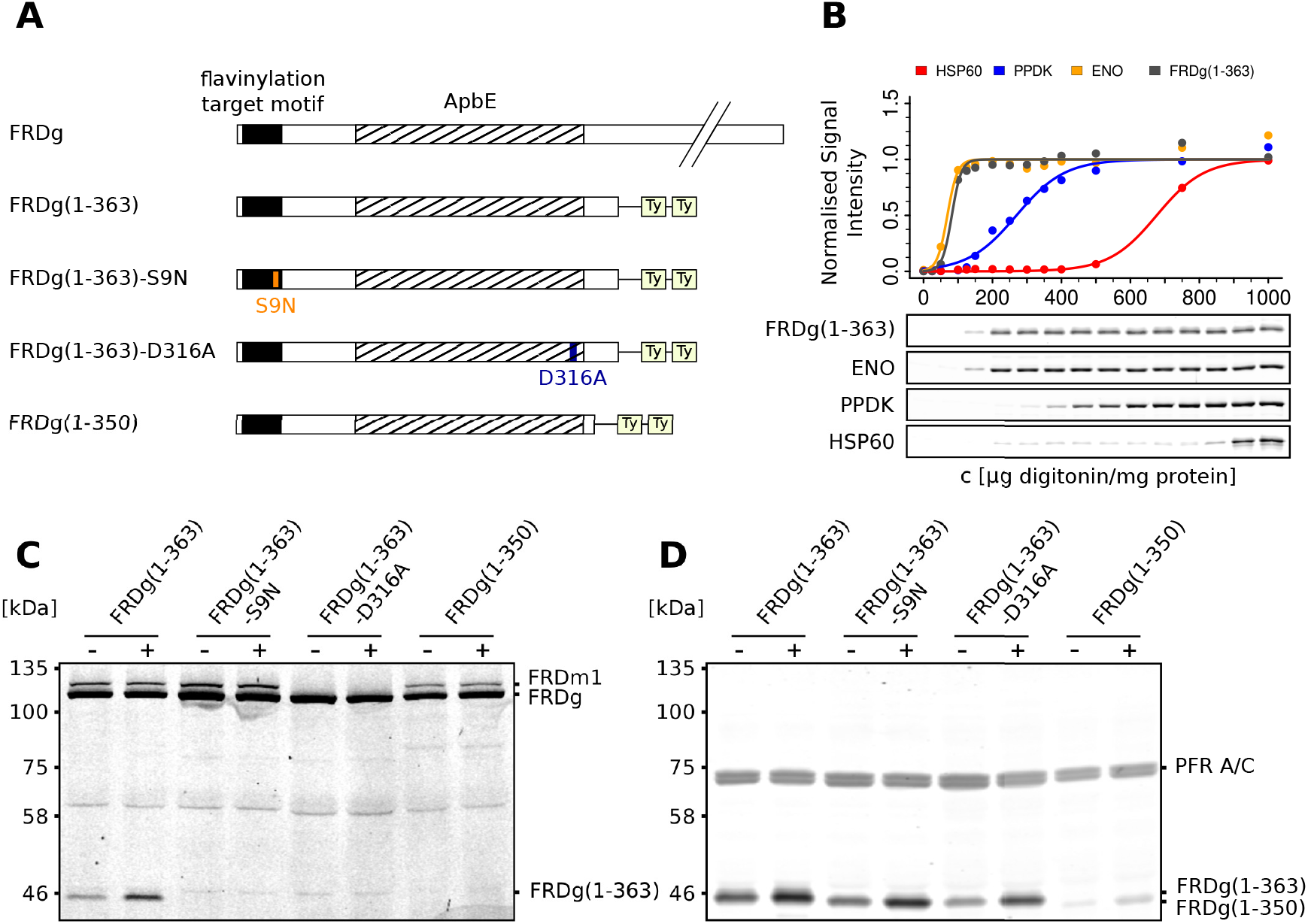
Flavinylation by the eukaryotic ApbE domain *in vivo*. **(A)** Schematic overview of transfected C-terminal TbFRDg truncation constructs, all Ty-tagged at the C-terminus. Wild type FRDg is shown as reference. The ApbE domain is indicated as striped box. The FTM is shown as black box; the Ser9 to Asn mutation is highlighted in orange, and the Asp316 to Ala mutation is marked in blue. The Ty epitopes are highlighted in light yellow. **(B)** Subcellular localisation of FRDg(1-363) by digitonin fractionation. Mitochondrial (*α*-HSP60), cytoplasmic (*α*-ENO) and glycosomal (*α*-PPDK) markers were compared to FRDg(1-363) (*α*-Ty1). Normalised signal intensities were plotted against the respective digitonin concentration. **(C)** In-gel fluorescence analysis for detection of covalently bound flavins in procyclic cell lines as indicated; (-) not induced, (+) tetracycline-induced. The FRDm1 band is not visible in the FRDg(1-363)-D316A cell line, as the parental AnTat1.1 90-13 clone showed declining expression of endogenous FRDm1, prior to this transfection. **(D)** Western blot corresponding to **C**. *α*-Ty1 was used for detection of FRDg truncations; *α*-PFR-A/C was used as loading control.

FRDg(1-363) encompassing the flavinylation target motif and the ApbE domain was covalently flavinylated (Figure 2C; band at 46 kDa). Replacement of Ser9, the flavin accepting residue in the orthologous *L. pyrrhocoris* FRD [32], with asparagine (FRDg(1-363)-S9N) removed all detectable flavinylation (Figure 2C). This confirms that Ser9 in the target motif DGRSSA<u>S</u> is the acceptor residue in *T. brucei*. The aspartic acid 284 of *Treponema pallidum* ApbE plays a major role in Mg^2+^ cofactor binding and is essential for flavinylation activity [62]. Comparison of the *T. brucei* FRDg N-terminus with the *L. pyrrhocoris* orthologue and the well-studied bacterial *T. pallidum* ApbE protein [62, 63] (Figure 1B) shows conservation of the ADALATA sequence at position 283-289 of *T. pallidum* ApbE. Site-directed mutagenesis of the residue homologous to Asp284 in *T. pallidum* ApbE was performed to obtain a catalytically dead ApbE domain (FRDg(1-363)-D316A). No flavinylation of this protein was detected albeit expressed at the same level as FRDg(1-363) (Figure 2C). Truncation of the ApbE domain from the C-terminal side in FRDg(1-350) also resulted in loss of flavinylation, probably by disrupting structural integrity close to the Mg^2+^-binding site. Lower abundance of FRDg(1-350) in four independent *T. brucei* cell lines, compared to FRDg(1-363), is compatible with reduced stability of this truncated protein (Figure 2D showing one representative line). These results led to the conclusion that the ApbE domain of FRDg has flavinylation activity and catalyses auto-flavinylation of the FRDg N-terminal motif. In addition, the activity does not depend on the glycosomal environment where FRDg is naturally localised, as FRDg(1-363) is expressed in the cytoplasm. Whether the auto-flavinylation proceeds as intramolecular or intermolecular reaction cannot be determined from the experiment (Figure 1C).

### 6.2 *Trans*-flavinylation of a 37 AA acceptor fragment is catalysed by the ApbE domain of FRDg

Next, we aimed at defining the minimal fragment sufficient to serve as flavinylation acceptor in a heterologous protein context. The first 37 amino acids of FRDg were fused to aconitase devoid of the N-terminal mitochondrial import signal (FRDg(1-37)-ACO; Figure 3A). Aconitase is a dual localisation enzyme in trypanosomes [40] and, upon removal of the mitochondrial import signal, is exclusively present in the cytoplasm. FRDg(1-37)-ACO was *tet*-inducibly expressed in the bloodstream form of *T. brucei* and its subcellular localisation verified by digitonin solubilisation (Figure 3B). As expected, no flavinylation of FRDg(1-37)-ACO was detected in absence of a functional ApbE protein in the same (cytoplasmic) compartment. Upon co-expression of FRDg(1-363), the FRDg(1-37)-ACO fusion protein became flavinylated, as evidenced by a fluorescent band at the expected size of the fusion protein (Figure 3C). Auto-flavinylation of FRDg(1-363) was at the same level compared to the control expressing FRDg(1-363) alone. We can draw four conclusions from these results: (1) there is no detectable, endogenous flavin transferase in the cytoplasm, (2) the first 37 amino acids of FRDg are sufficient to serve as flavinylation acceptor, (3) the predicted ApbE domain of FRDg is a functional flavin transferase, independently confirming evidence in Figure 2, (4) the ApbE domain of FRDg is able to *trans*-flavinylate a target motif *in vivo*, as reported for bacterial ApbEs.

**Figure 3:**
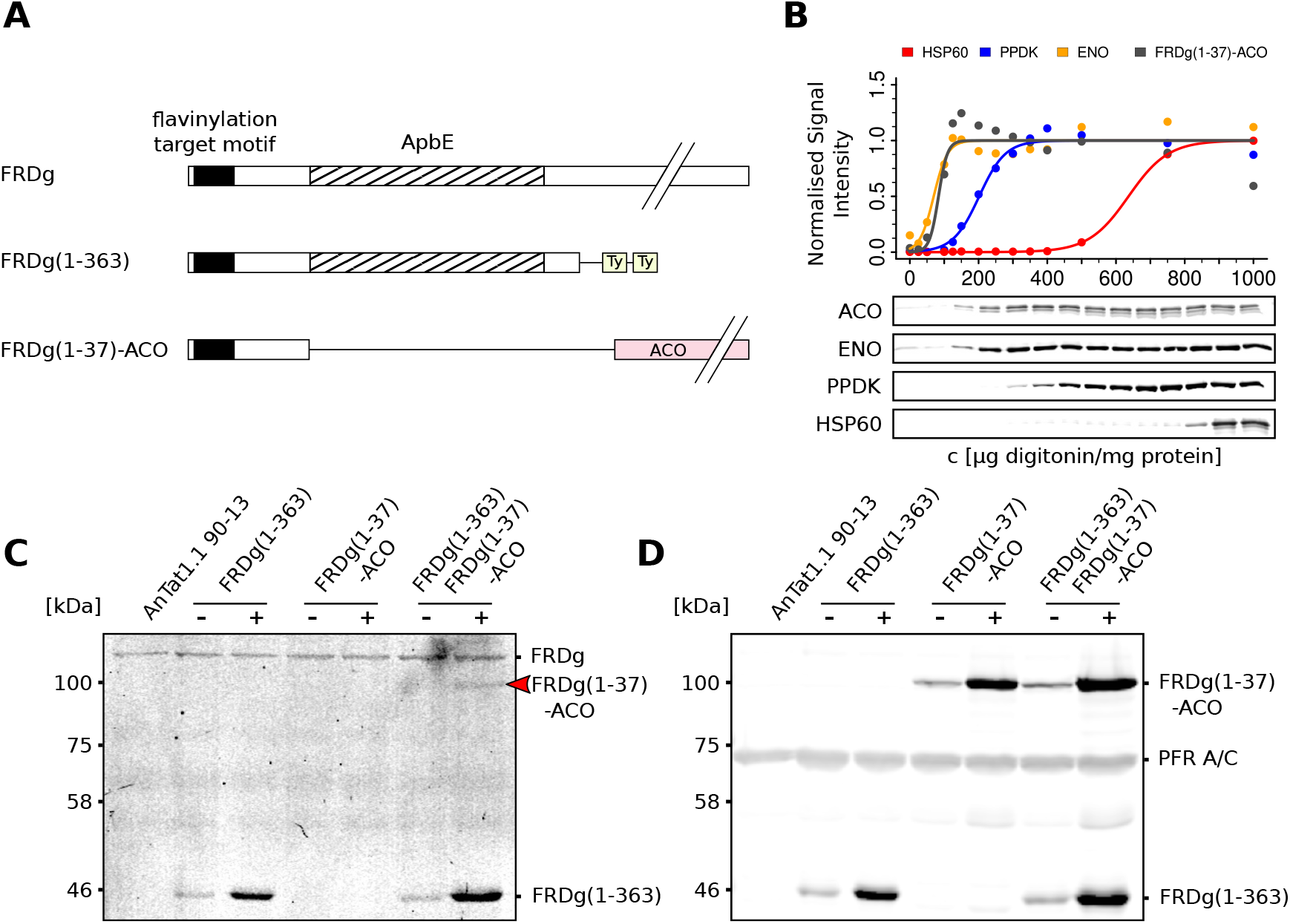
ApbE domain mediated *trans*-flavinylation of a small FTM-containing FRDg-peptide. **(A)** Schematic overview of transfected constructs FRDg(1-363) and FRDg(1-37) fused to cytoplasmic aconitase (FRDg(1-37)-ACO). FRDg(1-363) is Ty-tagged at the C-terminus. The aconitase coding region is highlighted in light red, the Ty epitopes in light yellow. **(B)** Digitonin fractionation showing cytoplasmic localisation of FRDg(1-37)-ACO in procyclic *T. brucei*. Rat *α*-ACO was used for detection of FRDg(1-37)-ACO, for marker proteins see legend to Figure 2B. **(C)** In-gel fluorescence for detection of covalently bound flavins in BSF *T. brucei* expressing FRDg(1-37)-ACO (marked by a red arrow) or FRDg(1-363) or both; (-) not induced, (+) tetracycline-induced. FRDm1 is expressed at levels below detection limit in BSF [29]. **(D)** Western blot corresponding to **C**. *α*-Ty1 was used for detection of FRDg(1-363), rabbit *α*-ACO for detection of FRDg(1-37)-ACO, and *α*-PFR-A/C was used as loading control.

### 6.3 Intramolecular cis-flavinylation of FRDg is more efficient

Since the ApbE domain in FRDg is able to flavinylate substrates *in trans* in the same way as bacterial ApbE enzymes, but is physically linked to its major substrate in the same polypeptide chain, alternative mechanisms of auto-flavinylation depicted in Figure 1C are possible: (1) *trans*-flavinylation of FRDg(1-363) by another FRDg(1-363) polypeptide or (2) intramolecular *cis*-flavinylation. We designed cell lines to compare *in vivo* the efficiency of both reactions. To test this in a physiologically relevant context, we fused the ApbE domain FRDg(1-363), its catalytically dead mutant FRDg(1-363)-D316A and N-terminal fragments only containing the flavinylation target motif to glycosomally localised isocitrate dehydrogenase (constructs FRDg(1-363)-IDHg, FRDg(1-363)-D316A-IDHg, FRDg(1-37)-IDHg, FRDg(1-9)-IDHg; Figure 4A). In the resulting transgenic cell lines, the glycosomal environment was maintained and *in vivo*-like concentration of enzyme and peptide substrate were ensured (Figure 4C). Targeting of all fusion proteins to the glycosome was verified by digitonin fractionation (Figure 4B, Figure S3G-I) and equivalent expression levels are documented in Figure 4D. The endogenous FRDg was present as internal reference and active flavin transferase in the glycosomes of all transgenic lines (Figure 4C). The flavinylation efficiency was expressed as ratio between covalent flavin attachment detected by direct in-gel fluorescence (Figure 4C) and the expression level of the fusion protein quantified by western blotting using an IDHg-specific antibody (Figure 4D). In addition, we normalised to the total amount of active flavin transferase in the glycosomes co-expressing FRDg(1-363)-IDHg and endogenous FRDg. The data from multiple independent experiments are displayed in Figure 4E and Figure S3A-F. The results clearly show a much stronger flavinylation of FRDg(1-363)-IDHg compared to the catalytically dead, but otherwise identical FRDg(1-363)-D316A-IDHg mutant that is unable to execute the intramolecular reaction (Figure 4C-E). We conclude that the intramolecular *cis*-flavinylation is 5x more efficient, this being a minimal estimate. The unusual fusion of the ApbE domain to its substrate FRD may provide a selective advantage during rapid metabolic adaptation of trypanosomes. The degree of *trans*-flavinylation of FRDg(1-37)-IDHg and FRDg(1-363)-D316A-IDHg by the endogenous FRDg is very similar, showing that the N-terminal 37 amino acids are fully sufficient as flavinylation target. This quantitatively confirms the independent results in Figure 3. In contrast, flavinylation of FRDg(1-9)-IDHg was not detected (Figure 4C). Apparently, the residues or the sequence environment C-terminal to the minimal consensus motif are also important.

**Figure 4:**
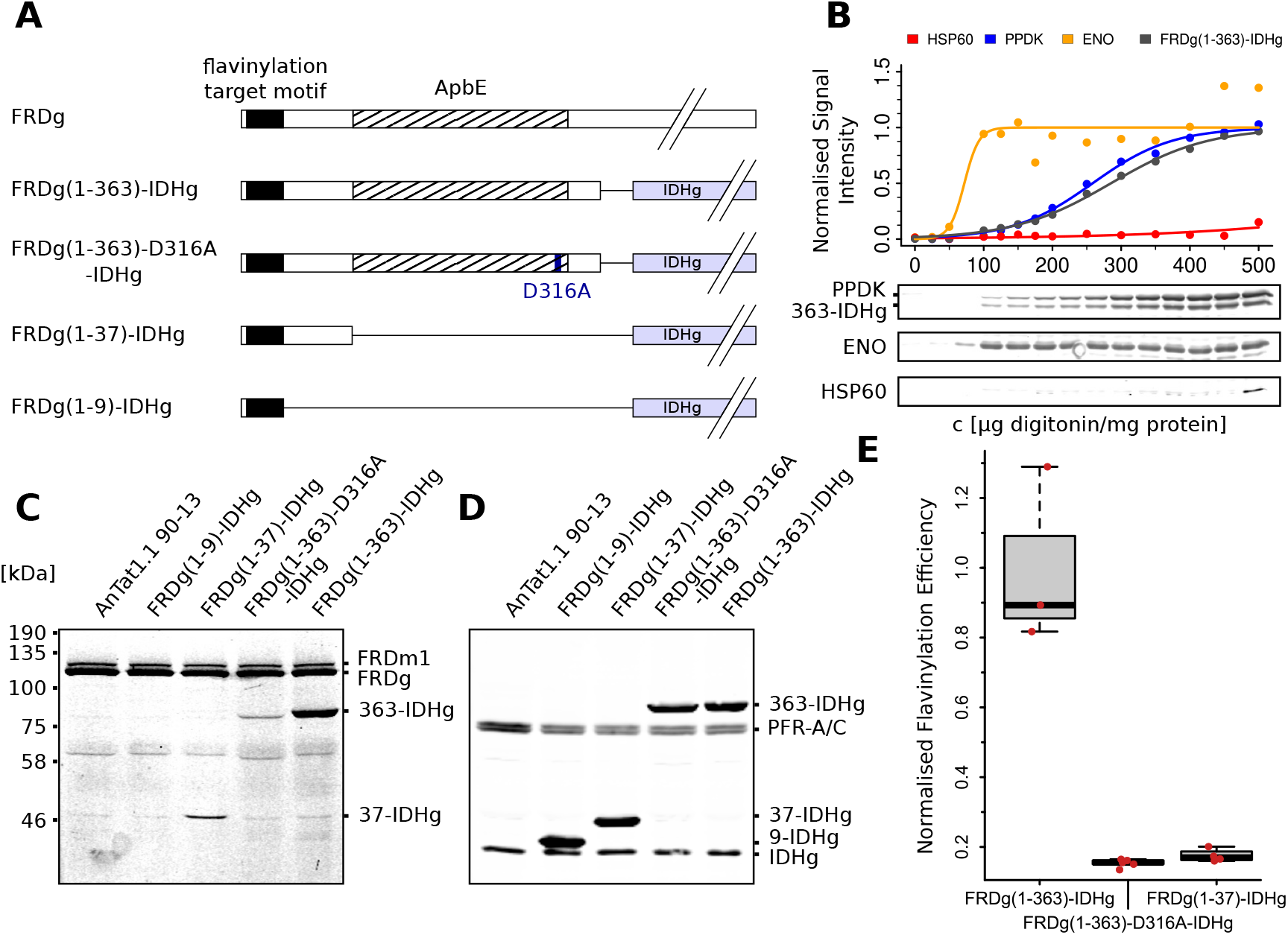
In glycosomes *cis*-flavinylation of FRDg is more efficient than *trans*-flavinylation. **(A)** Transfected IDHg-fusion constructs with IDHg coding region highlighted in light blue; see legend to Figure 2A. **(B)** Glycosomal localisation of FRDg(1-363)-IDHg verified by digitonin fractionation. *α*-IDHg was used for detection of the fusion protein, controls as detailed in legend to Figure 2B. **(C)** In-gel fluorescence for detection of covalently bound flavins in procyclic cell lines constitutively expressing the fusion proteins listed in **A**. **(D)** Western blot corresponding to the gel in **C**. *α*-IDHg was used for detection of the fusion proteins, and *α*-PFR-A/C was used as loading control. The following abbreviations were used in **C** and **D**: 9-IDHg for FRDg(1-9)-IDHg, 37-IDHg for FRDg(1-37)-IDHg, 363-IDHg for FRDg(1-363)-IDHg. **(E)** Quantification of flavinylation efficiency. The in-gel fluorescence signal of independent clonal cell lines (FRDg(1-363)-IDHg: n = 3; FRDg(1-363)-D316A-IDHg: n = 5; FRDg(1-37)-IDHg: n = 4; see Figure S3A-F) was normalised to the protein abundance as determined by western blotting of the same gel. The mean of the FRDg(1-363)-IDHg values was set as 100 %. In addition, we normalised the value for the FRDg(1-363)-IDHg cell line to the total amount of active flavin transferase in the glycosomes co-expressing FRDg(1-363)-IDHg and endogenous FRDg, using the flavin fluorescence signal. The standard box plot analysis shows median, first and third quartile of the resulting data, the whiskers define an outlier criterium at 1.5 times the interquartile range.

## 7 Discussion

This work shows that the ApbE-like domain present at the N-terminus of FRD of the kinetoplastid parasite *Trypanosoma brucei* has flavin transferase activity *in vivo*. ApbE-like domains are present in very few eukaryotic taxa and here functional activity of the predicted flavin transferase is documented for the first time to the best of our knowledge. To measure the activity *in vivo* we used transgenic expression in the cytoplasm, a subcellular compartment devoid of measurable flavin transferase activity, and quantified direct in-gel fluorescence of flavin covalently attached to the transgenic substrate. The specificity of the reaction was dually controlled by a catalytic dead mutant of the enzyme and by separate expression of the ApbE domain and the flavinylation target sequence. These controls also confirmed the absence of other covalent flavinylation activities in the cytoplasm of trypanosomes. The assay was performed in the bloodstream and procyclic developmental stages (Figure 2, Figure 3). Although the flavinylation target motif and the ApbE domain are juxtaposed in FRDg on the polypeptide chain, the ApbE domain alone is sufficient for *trans*-flavinylation of the substrate, as expected from the analogy to bacterial ApbE proteins (Figure 3).

The N-terminal 37 amino acids of FRDg that include the Dxx(s/t)(s/g)AS consensus motif [32] are sufficient as substrate for efficient flavinylation when fused to different proteins localised in the cytoplasm or in the glycosome. We also confirmed that Ser9 in this motif is essential for flavinylation [64], in agreement with Serebryakova *et al.* [32] who detected flavin attached to this site by MS in *L. pyrrhocoris*. The minimal motif fused to a heterologous protein (FRDg(1-9)-IDHg) was, however, not sufficient. Accordingly, the first 28 amino acids of FRDg in kinetoplastids are perfectly conserved [60] (Figure 1B) and secondary structures predicted upstream of the ApbE domain are preserved. This suggests additional structural or sequence requirements for recognition of the flavinylation target. In bacterial Na^+^-translocating NADH:quinone oxidoreductase [20], a conserved leucine is present close to the consensus motif at the position aligning to valine 11 of kinetoplastid FRDg (Figure 1B). Thus, a hydrophobic amino acid at this position might expand the minimally required motif.

Addressing the alternative mechanisms depicted in Figure 1C we found that *cis*-flavinylation of the FRDg target motif by the adjacent ApbE domain is at least 5-fold more efficient than *trans*-flavinylation. This conclusion follows from the large quantitative difference in flavinylation efficiency of the transfected intact FRDg N-terminus, compared to target proteins that only carry FRDg(1-37) or a catalytically dead, mutant ApbE domain FRDg(1-363)-D316A, all expressed in glycosomes in the presence of endogenous FRDg (Figure 4). Several arguments support validity of this conclusion: (1) all proteins were expressed at near 1:1 stoichiometry and at a level very close to endogenous FRDg in the native subcellular localisation in glycosomes, (2) analysis *in vivo* guarantees the physiological context, (3) the conclusion is independent of the endogenous flavinylation activity in glycosomes, that we assume to be exclusively FRDg, and (4) the assumption that full length FRDg and FRDg(1-363) have the same specific flavinylation activity is very likely correct, as the ApbE domain is structurally independent. The higher efficiency of the *cis*-flavinylation is consistent with single substrate Michael-Menten kinetics, compared to a random bi-substrate [65] mechanism for *trans*-flavinylation.

Flavoproteins have interesting properties for biotechnological applications. Flavin-based fluorescent proteins (FbFPs) are derived from light-oxygen voltage (LOV) sensing domains, utilised by plants and bacteria as blue-light photoreceptors [66, 67]. Their application as fluorescent reporters offers several advantages over GFP-like fluorescent proteins, such as oxygen-independent fluorescence [68, 69], a broad functional pH range (4-11) [70] and their small size – for iLOV approximately 10 kDa [71]. Other variants, such as miniSOG, can generate reactive singlet oxygen upon illumination, which is exploited for electron microscopy [72]. Furthermore, advances in signal intensity [73] allow application in super-resolution microscopy [74]. Known FbFPs use non-covalently bound flavin as cofactor, making them prone to denaturation by elevated temperatures [70]. Although variants with increased thermal stability exist [75], covalently bound flavin can tolerate even extended periods of thermal stress in the presence of detergents as shown by the assay used here for detection of flavinylation. We propose FRDg(1-37) as heat-stable, detergent-resistant fluorescent protein tag with a size of ~ 4 kDa, which is significantly smaller than a similar system proposed by Kang *et al.* [76] that utilises *Vibrio cholerae* Na^+^-translocating NADH:quinone oxidoreductase C (27 kDa) as tag in eukaryotic cells, which is flavinylated by ectopically expressed bacterial ApbE. The efficiency of the *trans*-flavinylation and sensitivity may limit general applicability as a tag for fluorescence microscopy, yet we envisage an application to analyse glycosomal import in kinetoplastids: the *trans*-flavinylation activity of endogenous glycosomally localised FRDg will be exploited to specifically detect and quantify any imported protein fused to the N-terminal FRDg(1-37) tag by direct in-gel fluorescence or flavin-specific antibodies [77] on denaturing PAGE. Only upon effective import, the tag on the fusion protein will be flavinylated *in trans* by FRDg, enabling analysis without prior cell fractionation. This N-terminal tagging approach is unlikely to interfere with the glycosomal import mechanism, which mostly relies on a C-terminal peroxisomal-targeting signal [78]. The strategy can be used to map glycosomal import sequence requirements, study the import mechanism and its regulation, or screen for novel glycosomal proteins.

Eukaryotic ApbE domains are rare and limited to few clades of the tree of life. The domain architecture of kinetoplastid FRDs (Figure 1A) is also predicted for some algal FRDs, such as *Symbiodinium microadriaticum* (GenBank accession: OLP85325), *Nannochloropsis gaditana* (GenBank accession: EWM27529) or *Nannochloropsis salina* (GenBank accession: TFJ85565). In contrast, the FRD of *Diplonema papillatum* (GenBank accession: LMZG01019705), that is in a clade close to kinetoplastids is not linked to an ApbE domain and no distinct ApbE protein could be detected in the *D. papillatum* genome so far. Independent fusion events of an ApbE domain to FRD in those phylogenetically distant taxa is compatible with an evolutionary advantage of intramolecular FRD flavinylation in certain physiological situations. The bacterial *K. pneumoniae* FRD provides a potential example of an evolutionary intermediate, where FRD and ApbE are transcribed from a single operon, but still form distinct proteins [19, 20].

Can we envisage a metabolic reason for the benefit of a more efficient *cis*-flavinylation of FRD in some organisms? Limited availability or transport of riboflavin [79] is a possibility, as trypanosomes are riboflavin auxotrophs. Most importantly, they depend on metabolic flexibility and adaptation to rapidly changing host environments in their parasitic life cycle. In the mammalian bloodstream, a simplified metabolism relies predominantly on the glucose supplied by the mammalian host. Pyruvate is the main glycolytic end-product [30, 80]. Upon transmission to the glucose-depleted, amino-acid-rich environment [81] in the tsetse fly insect vector, differentiation into the procyclic stage [82] results in major adaptions of the parasite’s metabolism. Energy sources are partially oxidised by aerobic fermentation to acetate or succinate [30]. In the glycosomes, this is achieved by establishment of the succinic branch of glycolysis [83], where FRDg regenerates NAD^+^ by reduction of fumarate to succinate [29, 30, 84]. To enable this rapid metabolic adaptation, glycosomes are remodelled depending on carbon source by degradation via pexophagy and *de novo* synthesis of glycosomes during differentiation of the parasite [85, 86]. The fusion of an ApbE flavin transferase domain to FRD may not only result in more efficient *cis*-flavinylation, as shown here (Figure 4E), but also in kinetically faster flavinylation of *de novo* synthesized FRDg. As flavinylation is a prerequisite for enzymatic activity [20], the kinetoplastid enzyme might be optimised for quickly re-establishing the glycosomal redox balance upon glycosome turnover and thereby provide an increased fitness in the rapidly changing environments the parasite inhabits. The same argument may be valid for the mitochondrial isoform FRDm1, as the trypanosomal mitochondrion undergoes extreme developmental restructuring in the life cycle.

## Supporting information

Graphical abstract

mini abstract/legend to graphical abstract

## 3 Abbreviations

ACO: aconitase
ApbE: alternative pyrimidine biosynthesis E
BSF: bloodstream form
ENO: enolase
FAD: flavin adenine dinucleotide
FbFP: flavin-based fluorescent protein
FMN: flavin mononucleotide
FRD: fumarate reductase
FRDg: glycosomal fumarate reductase
FRDm: mitochondrial fumarate reductase
FTM: flavinylation target motif
HSP60: heat shock protein 60
IDHg: glycosomal isocitrate dehydrogenase
LOV: light-oxygen voltage
PCF: procyclic form
PFR: paraflagellar rod
PPDK: pyruvate phosphate dikinase
tet: tetracycline

## 10 Acknowledgements

We thank D. Inaoka (NEKKEN, Nagasaki University) for critical reading of the manuscript and M. Wirth, I. Forné and A. Imhof (BMC, Ludwig-Maximilians-University Munich) for the mass spectrometry service.

## 12 Author contributions

RS, SB and MB designed research; RS performed research; RS, SB, FB and MB analysed data; RS and MB wrote the paper.

## 13 Supporting Information

**Figure S1:**
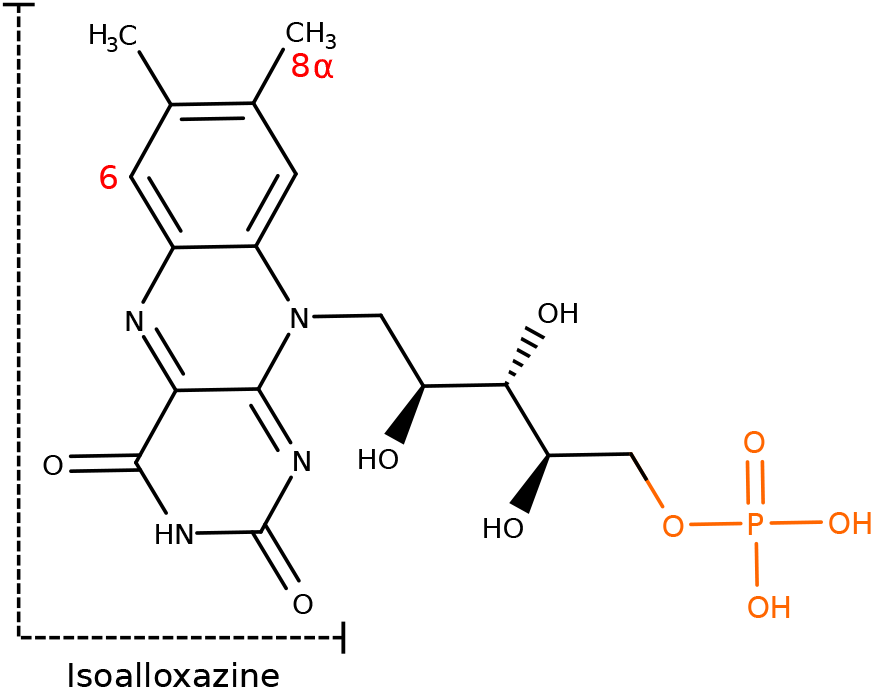
Structure of oxidised FMN. The C6 atom and the 8*α*-methyl group, which are subject to auto-catalytic protein attachment, are labelled in red. The phosphate group esterified by ApbE with the acceptor threonine or serine is highlighted in orange. MarvinSketch 20.13.0 (https://chemaxon.com/products/marvin) was used for chemical drawing.

**Figure S2:**
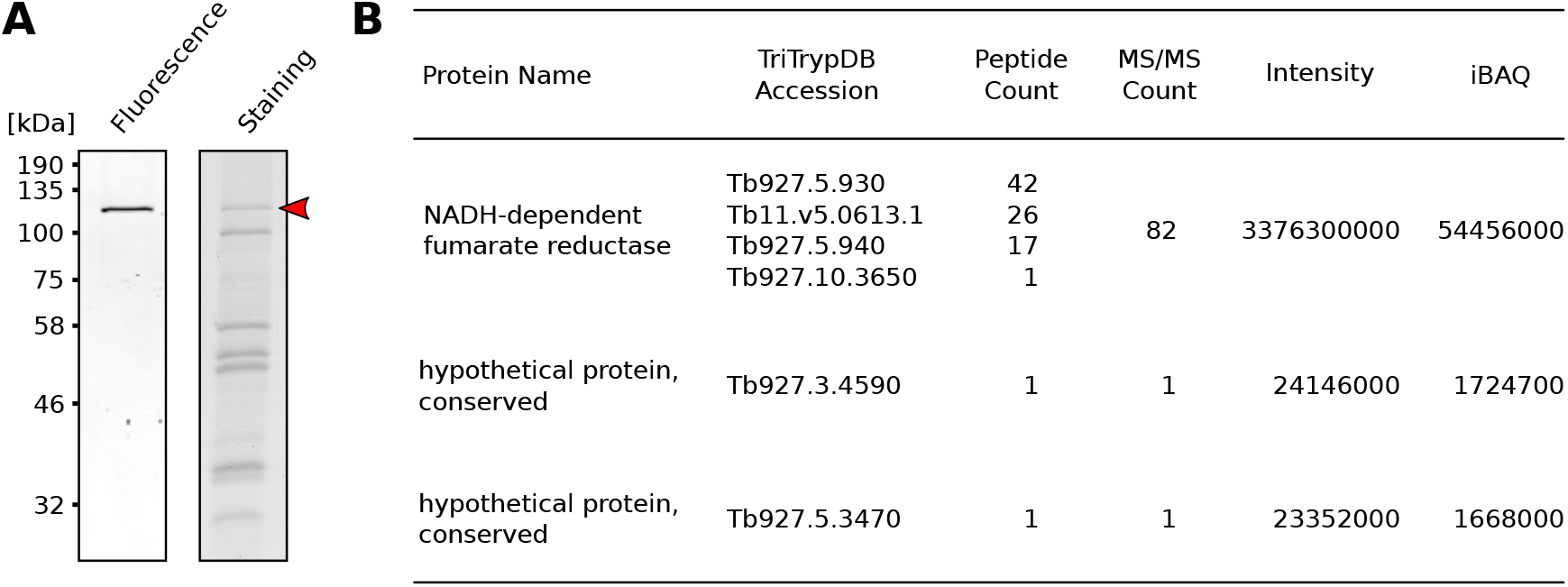
**(A)** In-gel fluorescence analysis (left) and matched Coomassie-staining (right) of a partially purified fraction of procyclic cell lysates. The excised band (marked by a red arrow) was destained, digested with trypsin and analysed by a LC MS/MS run and MALDI-TOF measurement. **(B)** Table of the top hits identified by mass spectrometry. MaxQuant [54] software was used.

**Figure S3:**
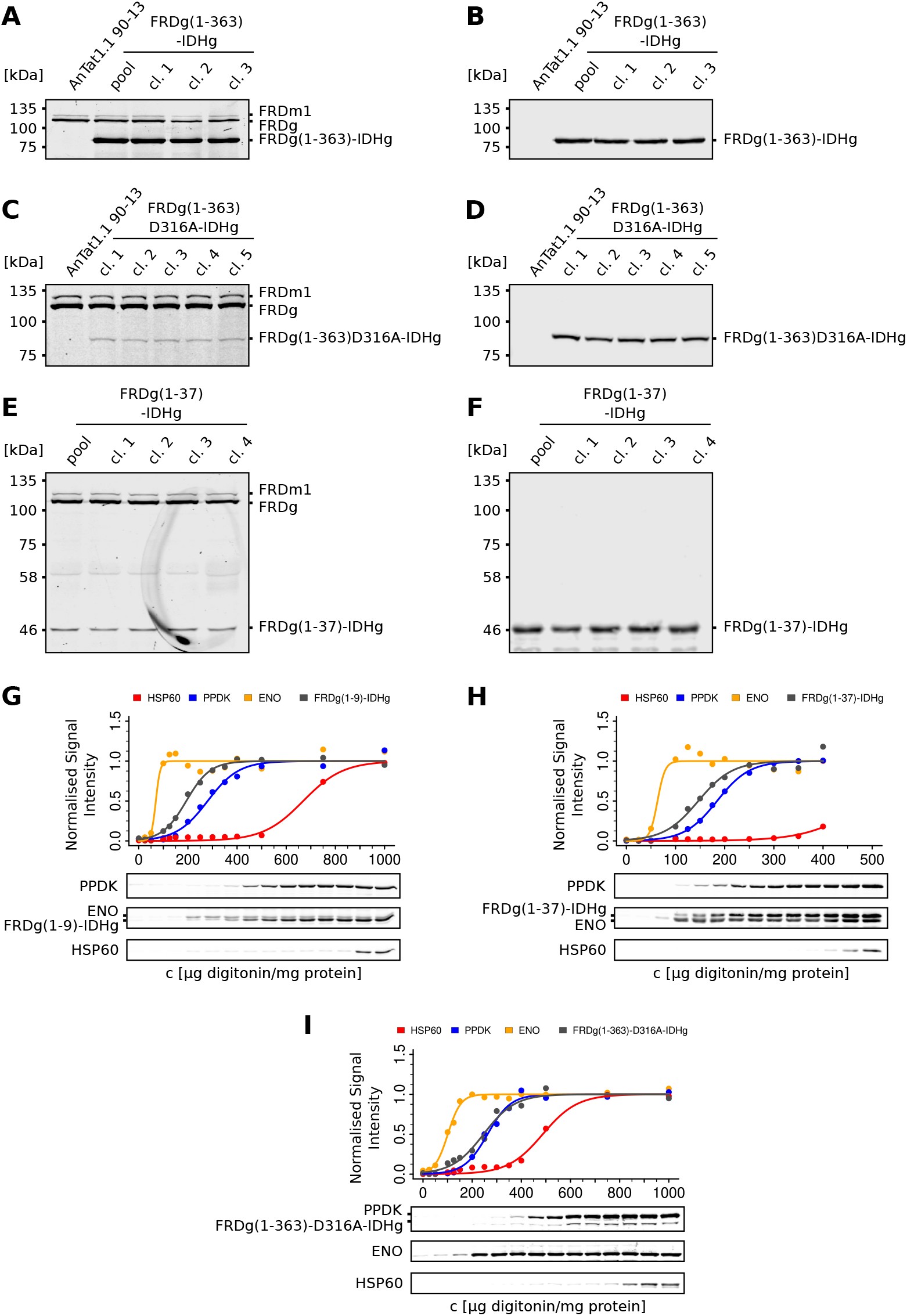
In-gel fluorescence and western blot analyses corresponding to the quantification shown in Figure 4E. **(A)** In-gel fluorescence analysis of one pool and three clonal (cl. 1-3) PCF cell lines constitutively expressing FRDg(1-363)-IDHg. **(C)** In-gel fluorescence analysis of five clonal (cl. 1-5) PCF cell lines constitutively expressing FRDg(1-363)-D316A-IDHg. **(E)** In-gel fluorescence analysis of one pool and four clonal (cl. 1-4) PCF cell lines constitutively expressing FRDg(1-37)-IDHg. **(B, D, F)** Western blots of the gels depicted in **A**, **C**, **E**, respectively; *α*-IDHg was used for detection. **(G, H, I)** Digitonin fractionations of FRDg(1-9)-IDHg (**G**), FRDg(1-37)-IDHg (**H**), and FRDg(1-363)-D316A-IDHg (**I**) were performed as described in legend to Figure 4. In **I** data points for digitonin concentrations *>* 400 *μg mg*^−1^ were omitted as outliers that interfered with the curve fitting algorithm.

