## Supplementary figures and images for "Efficient flavinylation of glycosomal fumarate reductase by its own ApbE domain in *Trypanosoma brucei*"

### Graphical abstract

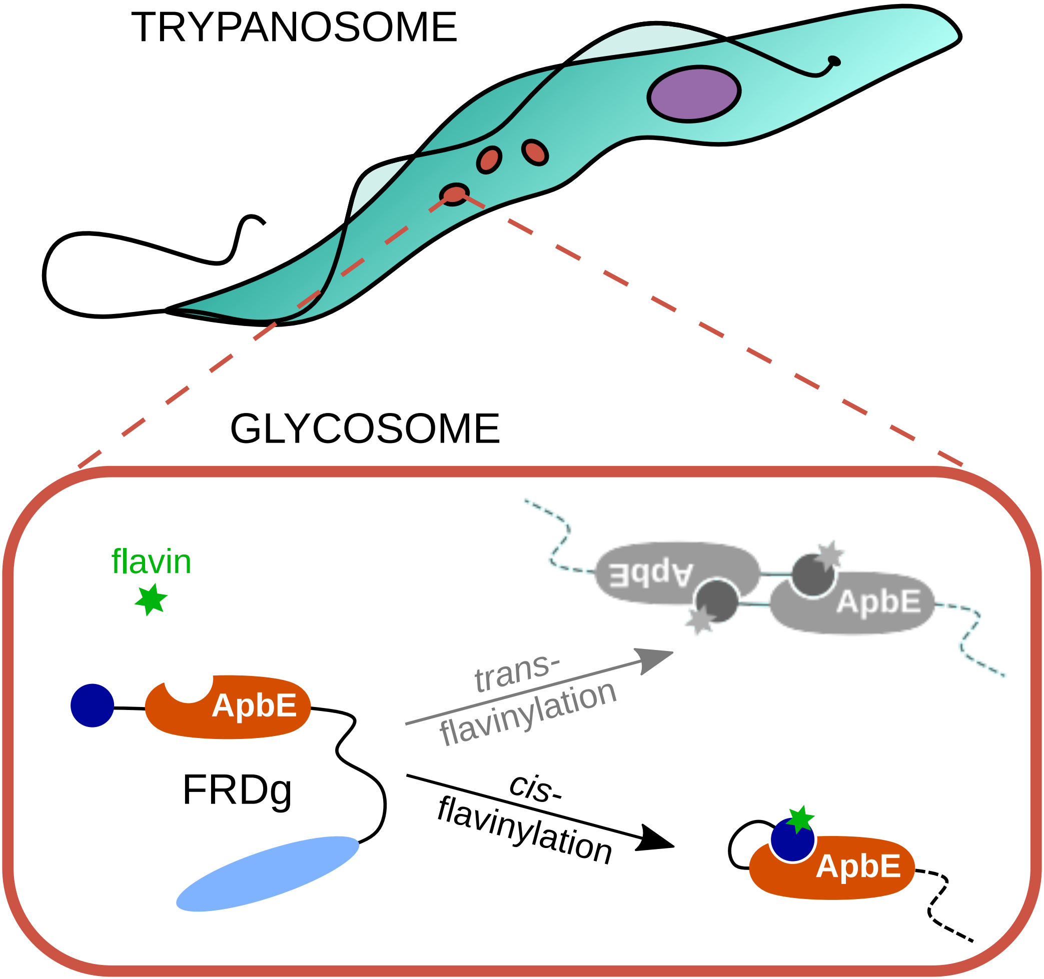
