## Supplementary material for "Efficient flavinylation of glycosomal fumarate reductase by its own ApbE domain in *Trypanosoma brucei*": mini abstract/legend to graphical abstract

In trypanosomes, a domain homologous to prokaryotic ApbE proteins (orange) that catalyse covalent protein flavinylation, is fused to fumarate reductase (FRD) adjacent to its flavinylation target site (blue). The domain can *trans*-flavinylate like prokaryotic ApbE, but intramolecular *cis*-flavinylation is more efficient and may provide a selective advantage by rapid FRD biogenesis. The N-terminal 37 amino acid flavinylation target of FRD can be used as conditional and heat stable fluorescence tag on heterologous proteins, a new tool to probe organelle import.
